# Analogs of the Dopamine Metabolite 5,6-Dihydroxyindole Bind Directly to and Activate the Nuclear Receptor Nurr1 (NR4A2)

**DOI:** 10.1101/2021.05.09.442997

**Authors:** Svetlana A. Kholodar, Geoffrey Lang, Wilian A. Cortopassi, Yoshie Iizuka, Harman S. Brah, Matthew P. Jacobson, Pamela M. England

## Abstract

The nuclear receptor-related 1 protein, Nurr1, is a transcription factor critical for the development and maintenance of dopamine-producing neurons in the substantia nigra pars compacta, a cell population that progressively loses the ability to make dopamine and degenerates in Parkinson’s disease. Recently, we demonstrated that Nurr1 binds directly to and is regulated by the endogenous dopamine metabolite 5,6-dihydroxyindole (DHI). Unfortunately, DHI is an unstable compound, and thus a poor tool for studying Nurr1 function. Here we report that 5-chloroindole, an unreactive analog of DHI, binds directly to the Nurr1 ligand binding domain with micromolar affinity and stimulates the activity of Nurr1, including the transcription of genes governing the synthesis and packaging of dopamine.

---

The nuclear receptor Nurr1 (NR4A2) plays critical roles in both developing and adult midbrain dopaminergic neurons, controlling the transcription of genes required for the synthesis (*TH)* and vesicular packaging (*VMAT2*) of dopamine, among other essential biological functions (e.g. management of oxidative stress, responsiveness to inflammatory signals).^1–3^ Clinical and animal data indicate that disrupted Nurr1 function contributes to inducing the dysregulation of dopaminergic neurons observed in the early stages of Parkinson’s disease (PD), as well as other dopamine-related CNS disorders (e.g. ALS, MS).^4–15^ Unraveling the complex biology of Nurr1 would be facilitated by bona fide Nurr1-targeting synthetic small molecules that can be used to directly modulate the receptor. Phenotypic assays have identified synthetic ligands that reportedly up-regulate transcription and protein levels of Nurr1 target genes, provide some degree of neuroprotection, and improve behavioral deficits in mouse models.^16–22^ However, there is little evidence that these compounds directly activate endogenous Nurr1, with the exception of the antimalarial drug amodiaquine and related analogs.^17, 23, 24^

Recently, we demonstrated that the endogenous dopamine metabolite 5,6-dihydroxyindole (DHI) stimulates the expression of *th* and *vmat2* in zebrafish and binds to the Nurr1 ligand binding domain (LBD) within a non-canonical ligand binding pocket^25^, forming a *reversible* covalent adduct with the side chain of Cys566, likely the result of a Michael addition to the oxidized, indolequinone (IQ) form of DHI (**Figure 1**, **Supplemental Figure 1A**).^26^ An endogenous prostaglandin (PGA1) has subsequently been shown to partially occupy this site, and form a covalent adduct with Cys5 66.^27^ DHI is unsuitable for robust biological studies, however, as it readily auto-oxidizes^28^ and polymerizes with itself and other molecules, in solution to form a chromogenic pigment and, in neurons to form neuromelanin.^29–31^ We, therefore, set out to identify unreactive analogs of DHI.

**Figure 1.**
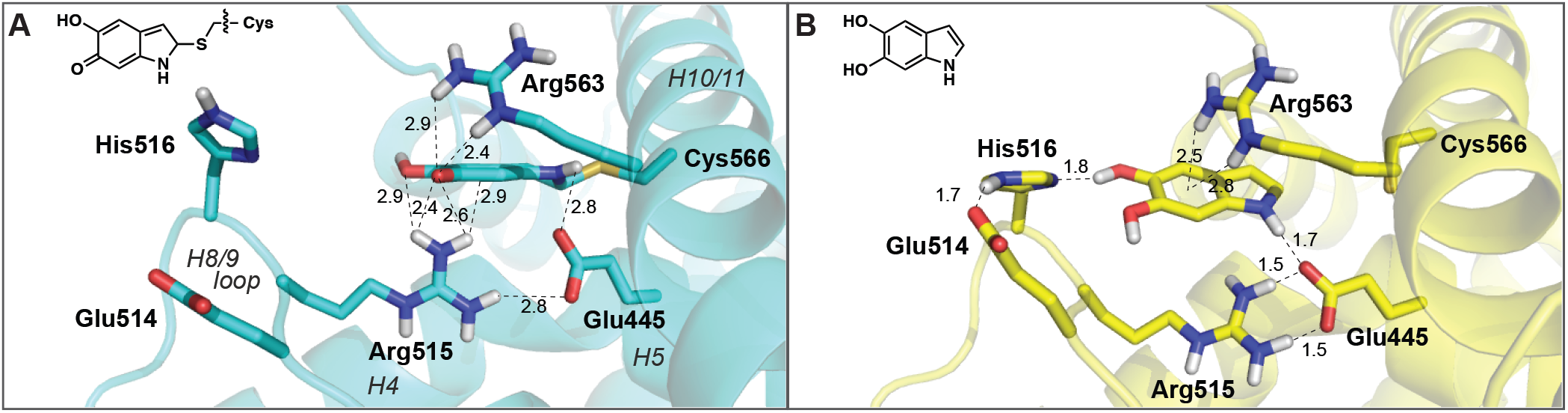
The binding of IQ and DHI to the Nurr1 LBD within the “566 site” is supported by a network molecular interactions. Close-up view of Nurr1 bound **(A)** covalently to indolequinone in the crystal structure (PDB:6DDA), and **(B)** non-covalently in a computational model of the unoxidized indole, DHI. In the QM/MM model, DHI is positioned farther away from H10/11 and tilted ~45 degrees along the plane of the ring, relative to the IQ, resulting in the formation of several new intra- and intermolecular interactions. Distances ≤3.0 Å only are shown here; see Supplemental Figure 1 for additional details. In the apo LBD structure (PDB: 1OVL), the guanidinium side chain of Arg563 is rotated ~180 degrees and forms an intramolecular bond with the carboxylate side chain of Glu445 (not shown).^32^

Previous biophysical and theoretical studies of indole (e.g. tryptophan analogs) interactions with cations established the importance of cation-π interactions in molecular recognition by proteins.^33, 34^ The crystal structure of the Nurr1-IQ complex shows a cation-π interaction between the side chain of Arg515 and the IQ adduct (**Figure 1A**). Weak electron density for the Arg563 side chain suggests that it is dynamic, and could form a second cation-π interaction with the ligand. Indeed, in quantum mechanical models of DHI bound non-covalently within the same “566 site,” a cation-π interaction with Arg563 appears important for stabilizing the aromatic indole system (**Figure 1B**). To identify unreactive analogs of DHI, we systematically replaced the 5- and 6-hydroxyl groups on the indole with a series of substituents expected to impact the strength of these cation-π interactions, and measured ligand affinities for the LBD in vitro, and activities in cells.

Approximately 20 substituted indoles, each predicted to bind within the LBD with poses nearly identical to DHI (**Supplemental Figure 1B**), were purchased from commercial suppliers and, for each compound, we determined (1) the molecular electrostatic potential (ESP) surface using density functional theory, (2) the affinity for the Nurr1 LBD using microscale thermophoresis (MST), (3) the activity against the full-length receptor using qPCR of Nurr1 target gene transcripts in MN9D^35^ cells, and (4) the cytotoxicity in MN9D cells. The affinity and activity across the entire series of compounds reveal that only indoles with a negative (red/orange) ESP surface, and thus most capable of forming a favorable cation-π interaction with the protein, exhibit saturable binding to the Nurr1 LBD (**Figure 2**, **Supplemental Figure 2A**). Dissociation constants could not be obtained for the amino- and hydroxyindoles, which we expected to bind to Nurr1, owing to their chemical instability in solution.^36–38^ Nonetheless, trends in the binding affinities among all of the halogenated indoles tested (*K_D_*: Br<Cl<<F) are consistent with previous reports surrounding the relative strength of cation-π interactions involving substituted indoles.^33, 34^ Intriguingly, the data also reveal that, among the indoles that bind to the LBD, only a subset also stimulate the transcription of Nurr1 target genes. In particular, whereas 5-chloro and 5-bromoindole bind with micromolar affinity (*K_D_* = 15 μM and 5 μM, respectively) and increase the expression of both *Th* (1.8- and 2.2-fold) and *Vmat2* (2.4- and 2.5-fold) in MN9D cells, the corresponding dihalogenated indoles bind with comparable affinity to the LBD, but do not modulate the expression of either gene. Cytotoxicity assays show that 5-chloroindole is not cytotoxic, whereas 5-bromoindole is among several indoles tested that are somewhat toxic to cells under certain conditions (≥10 μM, 24 h) (**Supplemental Figure 2C**).

**Figure 2.**
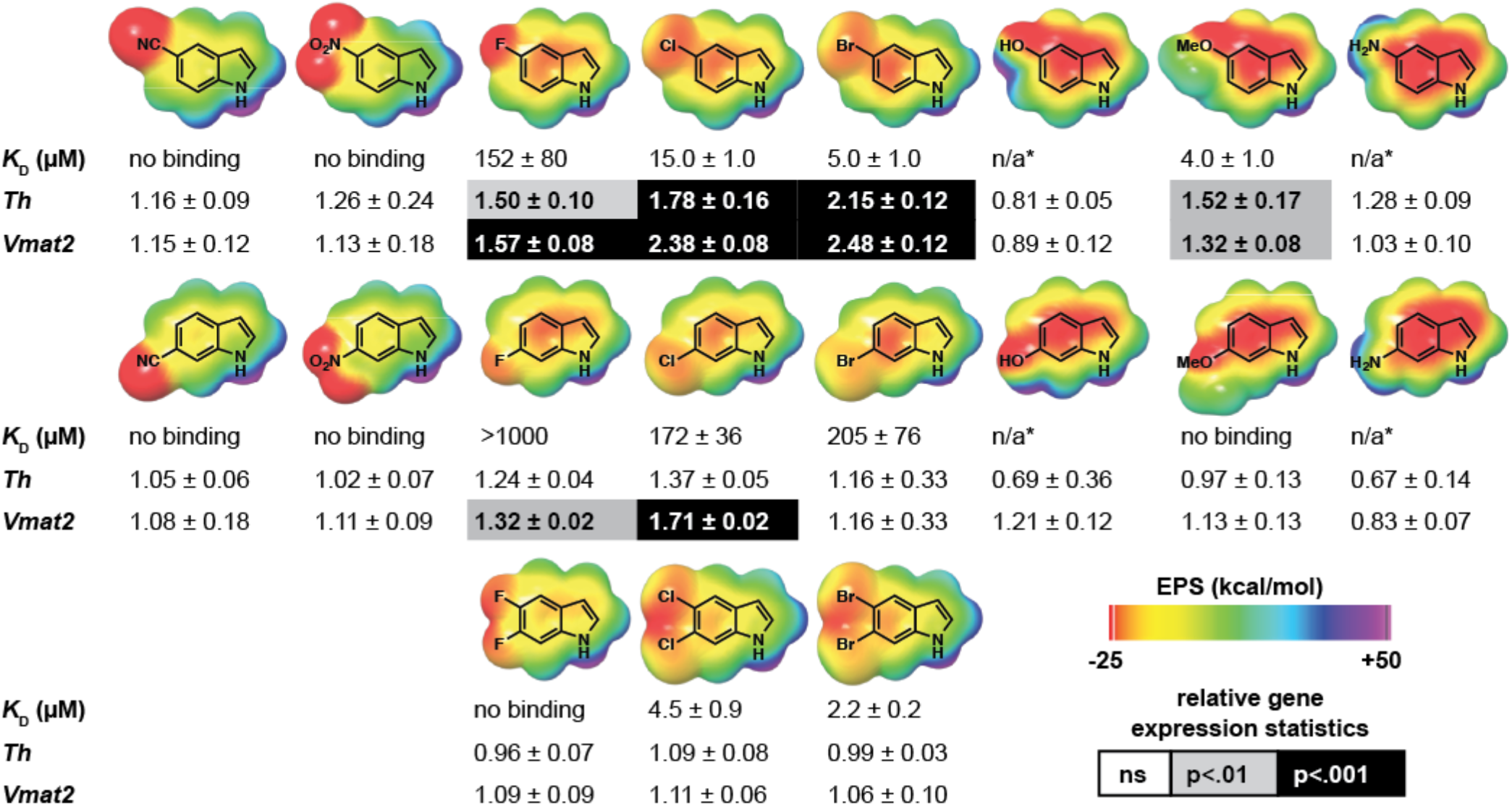
The DHI analogs 5-chloroindole and 5-bromoindole bind directly to stimulate the transcriptional activity of Nurr1. The molecular electrostatic potential (ESP) surface of each DHI analog was calculated using the 6-31G** basis sets, and bromine atoms treated with the LAV2P**. Binding affinity (*K_D_*) for the Nurr1 LBD was determined using microscale thermophoresis (MST). Relative expression levels of the Nurr1 target genes *Th* and *Vmat2* were determined using qPCR analyses of mRNA isolated from MN9D cells following treatment with each compound (10 μM, 24 h). Transcript levels for each target gene were normalized to the housekeeping gene *Hprt*, and reported as the fold-change relative to cells treated with vehicle (DMSO) only. All experimental values are the result of three or more independent measurements ± SD. Detailed experimental protocols are described in Supporting Information *Note: data acquisition for these compounds is precluded by their chemical instability (see references 36-38). In particular, robust binding data could not be obtained because of initial fluorescence quenching and photobleaching (5- and 6-aminoindole), and autooxidation and polymerization in solution (5- and 6-hydroxyindole, 5- and 6-aminoindole). These compounds also exhibited significant cytotoxicity (see Supplemental Figure 2C).

Several experiments with 5-chloroindole support that the observed binding affinity and effect on gene transcription are due to direct interaction of the small molecule with Nurr1 (**Supplemental Figure 3**). First, increasing concentrations of the surfactant used in the MST binding assay only has a minor effect on the affinity of 5-chloroindole for the Nurr1 LBD (**Supplemental Figure 3A**), consistent with the values being due to individual molecules binding specifically to the protein, as opposed to aggregated indoles driving the response through non-specific interactions. Second, 5-chloroindole stimulates the activity of Nurr1 in two different luciferase reporter assays, one utilizing a chimeric Nurr1-LBD_Gal4DBD protein that binds to the Gal4 response element to drive luciferase expression, the other relying on binding of the fulllength receptor to the NBRE response element (**Supplemental Figure 3B**). Third, the stimulatory effect of 5-chloroindole on the expression of dopamine-related target genes in MN9D cells is inhibited by siRNA specific for Nurr1 (**Supplemental Figure 3C**). Lastly, 5-chloroindole does not exhibit saturable binding to the LBD of RXRa (**Supplemental Figure 3C**), demonstrating the effect on gene transcription is *not* due to ligand binding to RXRa within an Nurr1-RXRα heterodimer.

To investigate the molecular basis for the intriguing difference in activity between the 5-versus 5,6-halogenated indoles, we mutated residues in the 566 site postulated to provide stabilizing interactions with these indoles. In both the x-ray structure of the Cys566-IQ adduct (vide supra) and our QM/MM models of DHI non-covalently bound, cation-π interactions are important for stabilizing the ligand (**Figure 1**). In the QM/MM model, the side chain of Arg563 participates in the cation-π interaction. In addition, a halogen bond forms between the side chain of His516 and the halogen at the 5-positon (**Supplemental Figure 1C)**. Using MST, we characterized the binding affinity of the four indoles for the Arg563Ala and His516Ala single and double mutant proteins, with close attention paid to changes in the response amplitude, a metric that is very sensitive to changes in the size, charge and solvation shell of the protein, and thus a reporter of differences in protein conformation.^39^

The binding of 5-bromoindole to the Arg563Ala mutant Nurr1 LBD, alone or in combination with the His516Ala mutation, is eliminated, consistent with loss of a stabilizing cation-π interaction. The MST response amplitudes fall below the limit of robust signal detection for both mutants, and are significantly different from the large response amplitude observed for binding to the wildtype protein. Similarly, the response amplitude associated with 5-bromoindole binding to the His516Ala mutant is significantly different from that of the wildtype, though the binding affinity is statistically unchanged. The precise roles histidine side chains play in ligand-receptor interactions depend on the pKa of the side chain, which is influenced by the local environment inclusive of the ligand itself, making it generally difficult to predict the nature of the interaction.^40–43^ The affinity of 5-chloroindole for each of the mutants is *improved* (~2-fold) relative to the wildtype protein, whereas the response amplitudes are markedly different. In striking contrast, the affinities of the corresponding dihalogenated indoles for each of the mutant proteins are reduced (~3-7-fold), while the response amplitudes are not significantly different from the wildtype protein. Control assays suggest it is unlikely that changes in protein stability account for the differences in binding affinity between the wildtype and mutant proteins. The Arg563Ala mutation reduces the protein stability by ~3.6°, likely because the guanidinium side chain forms a hydrogen bond with the carboxylate side chain of Glu445 in the unliganded structure (PDB:1OVL), and the His516Ala mutation slightly increases the thermal stability (~0.4°) of the LBD (**Supplemental Figure 5**).

While the observed changes in binding affinity are equivocal, the changes in MST response amplitude between the wildtype and mutant proteins, combined with the effects of the 5-versus 5,6-substitued indoles on target gene transcription, point to a model in which there are two (or more) indole binding sites within the Nurr1 LBD. Binding of 5-chloro and 5-bromoindole to the 566 site supports the transcription of target genes *Th* and *Vmat2*, whereas binding of 5,6-dichloro and 5,6-dibromoindole to the other site(s) does not. The notion that there is more than one binding site for indoles within the Nurr1 LBD is in agreement with our previous study detailing the binding of DHI to Nurr1.^26^ As well, biophysical and computational studies of the interaction of bis-indoles and other compounds targeting Nurr1 also suggest the presence of additional binding sites within the LBD.^24, 44, 45^ Alternatively, it is possible these indoles bind to the same site, with the 5-substituted indoles inducing markedly different changes in the protein structure (and potentially interactions with co-regulatory proteins associated with transcription) than the corresponding 5,6-disubstitued indoles.

In conclusion, we have demonstrated that 5-chloroindole, a non-cytotoxic stable analog of the dopamine metabolite DHI, is suitable for directly probing the structure and function of Nurr1 (**Figure 3**). The affinity of 5-chloroindole for the Nurr1 LBD is comparable to DHI and amodiaquine in vitro and in luciferase reporter assays, whereas the potency in MN9D cells is superior to that of DHI and amodiaquine. Future studies will be aimed at discovering target genes possibly regulated by the 5,6-dihalogenated indoles and structure-activity relationship studies associated with different substitution patterns around the indole ring.

**Figure 3.**
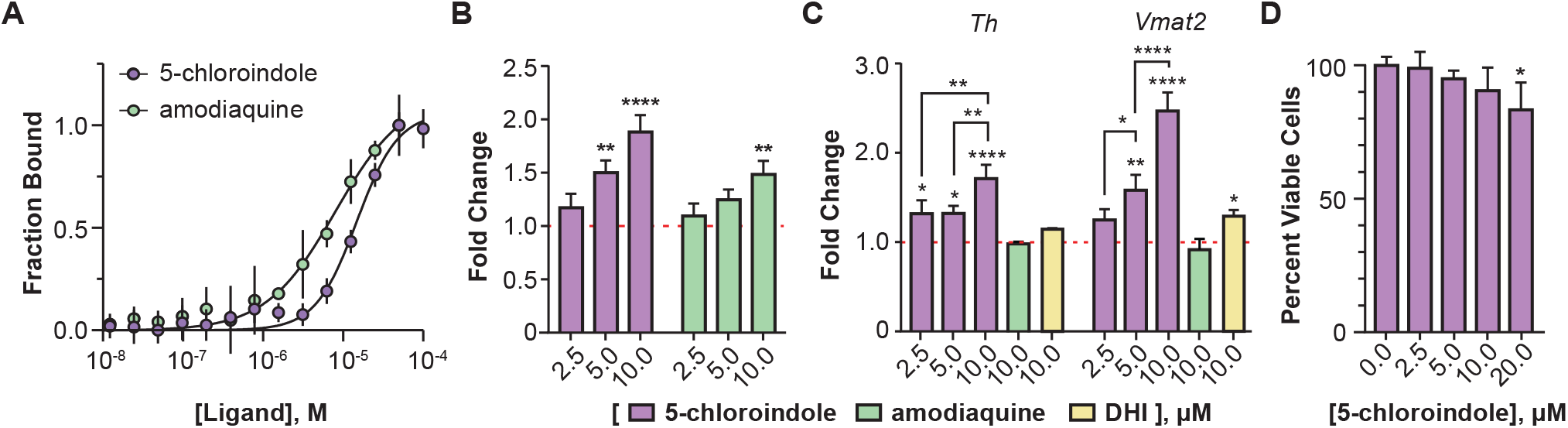
The DHI analog 5-chloroindole binds directly to and stimulates the activity of Nurr1. **(A)** Binding affinity of 5-chloroindole (15.0 ± 1.2 μM) and amodiaquine (8.0 ± 1.5 μM) measured by MST with 0.1% of the surfactant Pluronic F127. **(B)** Concentration-dependent stimulatory effect of 5-chloroindole and amodiaquine on the activity of a chimeric Nurr1-LBD_Gal4DBD protein to produce luciferase (6 h treatment). **(C)** Concentration dependent effect of 5-chloroindole on endogenous full-length Nurr1 to upregulate expression of *Th* and *Vmat2* in MN9D cells (24 h treatment) and effect of DHI and amodiaquine at 10 μM (24 h). Fold changes are relative to vehicle (DMSO) only. **(D)** Effect of 5-chloroindole on the viability of MN9D cells following treatment for 24 h. All experimental values are the result of three or more independent biological replicates ± standard deviation, with *p<0.05, **p<0.01, ***p<0.001, ****p<0.0001 by one-way ANOVA, in comparison with the response for vehicle treatment (DMSO).

## Supporting information

Kholodar et al., Supporting Information

## Supporting Information

Experimental Protocols and Supplementary Figures are included in supporting information. This material is available free of charge on the ACS Publications website.

## Acknowledgement

This work was supported by grants from the NIH (1R01NS108404-01) and the Program for Breakthrough Biomedical Research that is partially funded by the Sandler Foundation. We thank Dr. Thomas Perlmann (Karolinska Institute) and Dr. Jacques Drouin (Institut de Recherches Cliniques de Montréal, Canada) for generously providing us with the MN9D Tet-ON cell line and the NBREx3-POMC-Luc reporter plasmid, respectively.

