## Supplementary material for "Analogs of the Dopamine Metabolite 5,6-Dihydroxyindole Bind Directly to and Activate the Nuclear Receptor Nurr1 (NR4A2)": Kholodar et al., Supporting Information

Svetlana A. Kholodar, Geoffrey Lang, Wilian A. Cortopassi, Yoshie Iizuka, Harman S. Brah, Matthew P. Jacobson, Pamela M. England*
Departments of Pharmaceutical Chemistry and Cellular Molecular Pharmacology, University of California San Francisco, San Francisco, CA 95158, United States.

**Supporting Information**

### SUPPLEMENTAL FIGURES

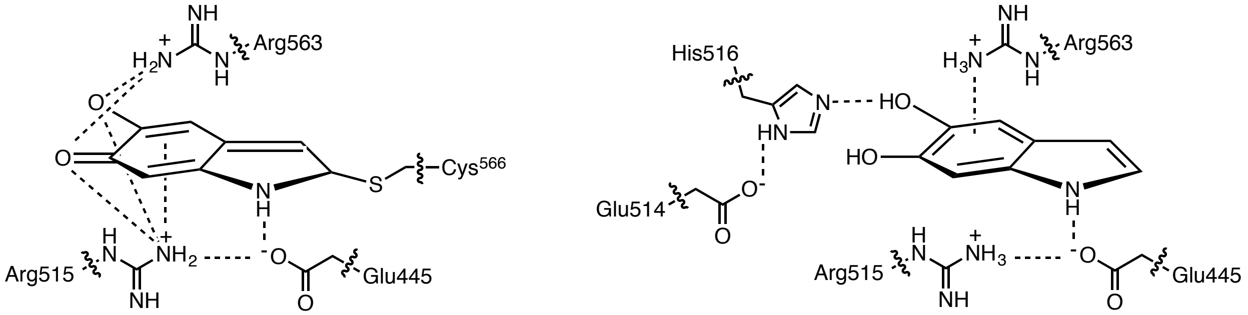


| Residue/Interaction | Interaction | IQ (X-ray) | DHI (QM/MM) | 5-Chloroindole (QM/MM) |
|---|---|---|---|---|
| Glu445-NH(ligand, pyrole) | Hydrogen bond | 2.8 | 1.7 | 1.7 |
| Glu445-Arg515 | Ionic salt bridge | 2.8, 3.3 | 1.5, 1.5 | 1.5, 1.6 |
| Glu514-His516(HE2) | Hydrogen bond | 6.5 | 1.7 | 1.7 |
| His516(ND1)-5-HO(ligand) | Hydrogen bond | 6.1 | 1.8 | NA |
| Arg515-5-oxo(ligand), 5-Cl(ligand) | Hydrogen bond | 2.9 | 5.8 | 6.0 |
| Arg515-6-oxo(ligand) | Hydrogen bond | 2.4, 2.6 | 5.0, 5.7 | NA |
| Arg563-5-oxo(ligand), 5-Cl(ligand) | Hydrogen bond | 2.9 | 3.6 | 3.6 |
| Arg563-6-oxo(ligand) | Hydrogen bond | 2.4, 2.9 | 4.1 | NA |
| His516(ND1)-5-Cl | Halogen bond | NA | NA | 3.0 |
| Arg515-PhCtr | Cation-π | 2.9 | 3.8 | 3.6, 4.2 |
| Arg563-PhCtr | Cation-π | 3.8, 3.9 | 2.5, 2.8 | 2.6, 2.7 |

**Supplemental Figure 1A.** The binding of indoles to the Nurr1 LBD is stabilized by networks of hydrogen, halogen, cation-π, and ionic bonds. **(Top)** Chemical structures showing interactions between amino acid side chains within the Nurr1 LBD and bound ligands; only interactions with distances ≤3.0 Å are shown. **(Bottom)** Table showing the physical distances in Å between amino acid side chains the bound ligands. Distances ≤3.0 Å are shown in black and distances >3 Å are shown in grey.

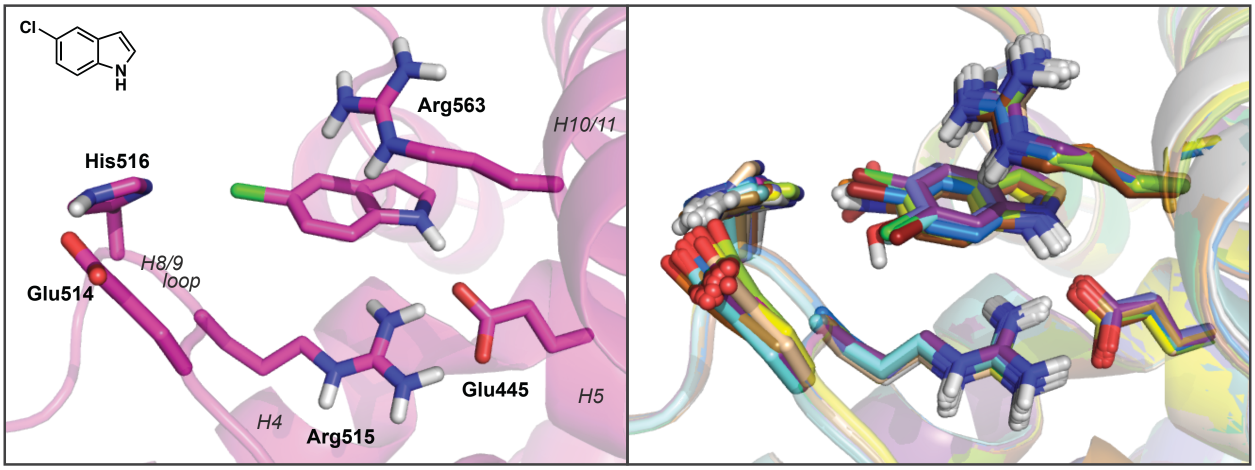


**Supplemental Figure 1B**. Substituted indoles are predicted to bind with nearly identical poses to Nurr1 in computational (QM/MM) models. **(Left)** Model of 5-chloroindole bound to Nurr1. **(Right)** Overlay of computational models for all of the halogenated, and 5-substitued, indoles evaluated in the present study.

| | Modeled Interaction with His516 | His516 pKa | Single Point Interaction Energy | Ranking (ΔG, gas phase) | Measured Affinity ($K_D$) |
|---|---|---|---|---|---|
| 5-bromoindole 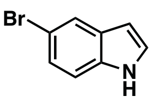  | 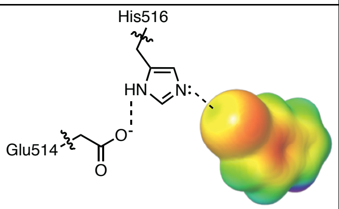  | 7.1 | -1.3 | 4 | 5.0 ± 1.0 μM |
| 5-chloroindole 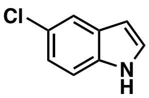  | 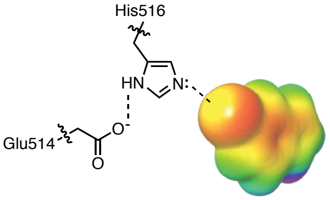  | 7.0 | 0 | 6 | 15.0 ± 1.0 |
| 5-fluoroindole 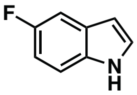  | 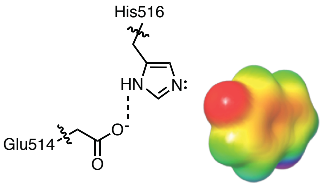  | 7.8 | 5 | 8 | 152 ± 80 μM |

**Supplemental Figure 1C**. Binding of 5-bromo- and 5-chloroindole to Nurr1 is predicted to be stabilized by a halogen bond with His516. Lateral views of the molecular ESP surfaces for the 5-halogen-substituted indoles highlight the interaction between the lone pair of electrons on His516 and the sigma hole within the bromo and chloro substituents. The deficiency in electron density in the outer lobe of the $p_z$ orbital of 5-bromoindole and 5-chloroindole results in a relatively more positive electrostatic potential surface in this region, compared to 5-fluoroindole. The relative pKa values, interaction energies, and measured binding affinities are consistent with the proposed halogen bond between His516 and a subset of the halogenated indoles. The pKa values were predicted using propKa 3.1 after QM/MM optimization of the non-covalently bound indoles. The single point interaction energies were calculated with the LMP2/cc-pVDZ** level of theory in the gas phase. The coordinates of the complexes were taken from the QM/MM optimized structures at the DFT-D3/LACVP* level of theory. Ranking is among all of the 5-substituted indoles in the present study.

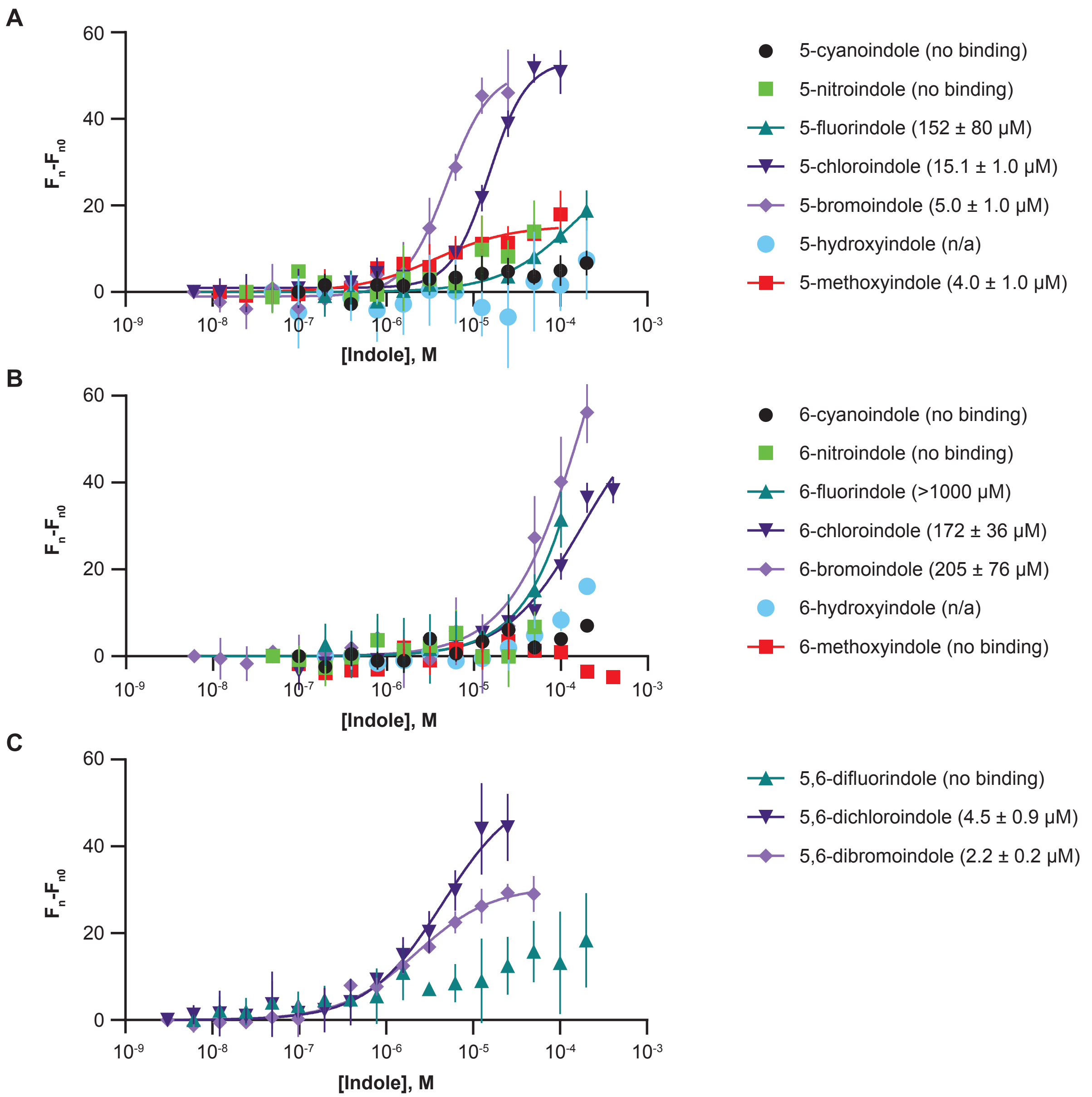


**Supplemental Figure 2A.** A subset of the halogenated indoles bind to the Nurr1 ligand binding domain. Microscale thermophoresis (MST) binding isotherms for **(A)** 5-substituted, **(B)** 6-substituted, and **(C)** 5,6-dihalogenated indoles and the Nurr1 LBD are obtained by plotting the change in thermophoresis ($F_n$-$F_{n0}$) versus the concentration of the compound tested ([Indole], M). All experimental values are the result of three or more independent measurements ± SD. All data were best fit to a single site, except for 5-chloro and 5-bromoindole, which required used of the Hill equation. Note: The Hill coefficient ($n_H$) for 5-chloroindole (1.9 ± 0.2) and 5-bromoindole (1.9 ± 0.3) are both >1, whereas the value for all other compounds is unity within the error. A Hill coefficient greater than one typically indicates cooperative binding of ligands, with the absolute value setting the lower limit for the number of interacting binding sites (see Weiss, J. N. The Hill equation revisited: uses and misuses, *FASEB J. 11*, 835-841, 1997). However, we observed a significant change in the Hill coefficient with increasing concentrations of surfactant for 5-chloroindole (see Supplemental Figure 3D), possibly due to partial denaturation of the protein and concomitant loss of one of the indole binding sites. Alternatively, increasing concentrations of surfactant may have broken up compound nanoaggregates that falsely signaled cooperative binding of two indoles.

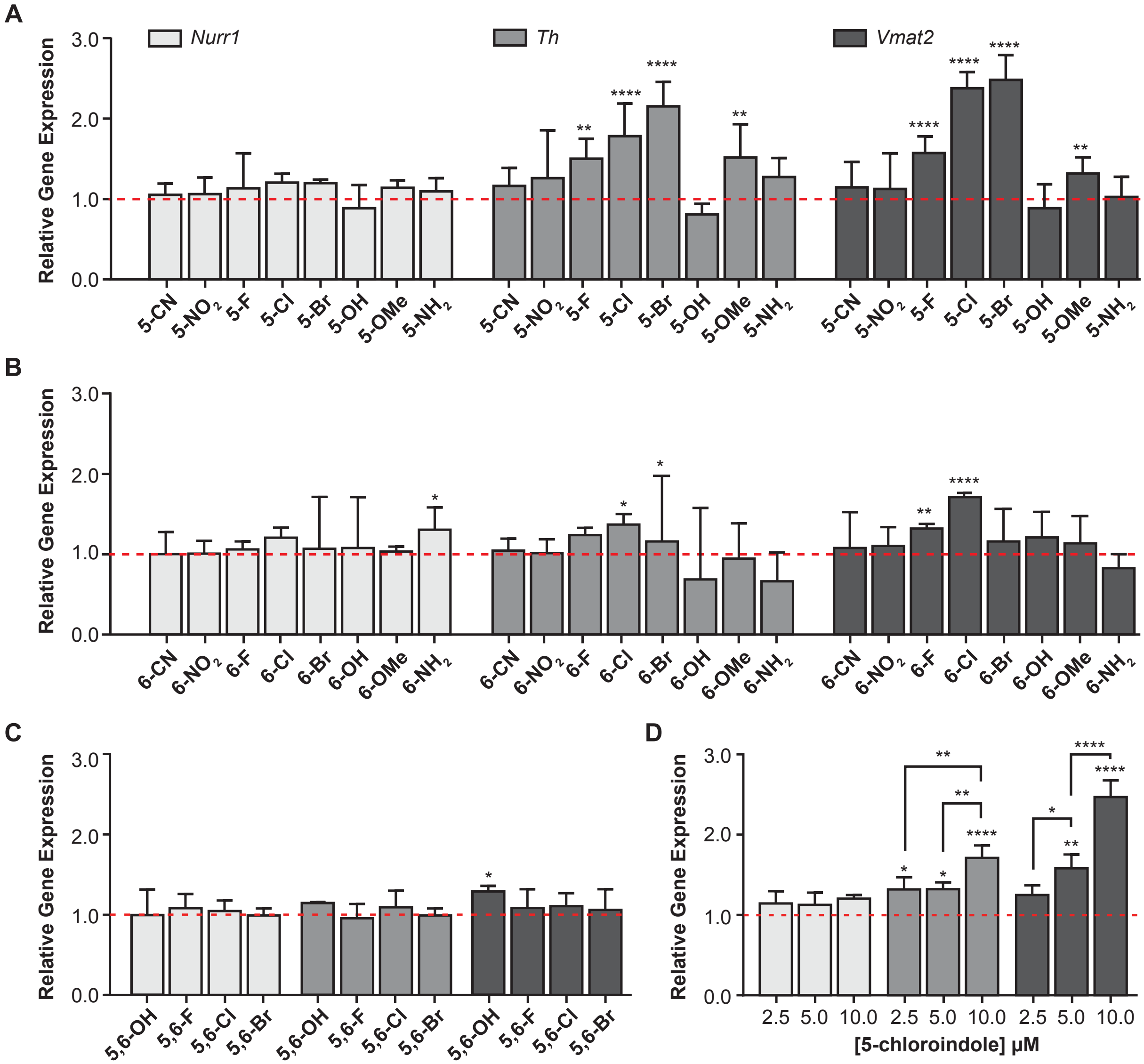


**Supplemental Figure 2B.** Only a subset of the indoles that bind to Nurr1 also stimulate the transcription of Nurr1 target genes in MN9D cells. The effect of **(A)** 5-substituted, **(B)** 6-substituted, and **(C)** 5,6-dihalogenated indoles (10 μM, 24 h) relative to vehicle (DMSO) only (dashed red line) on the expression of *Nurr1*, *Th*, and *Vmat2* was quantified by qPCR as described in Supplemental Information. **(D)** The effects of 5-chloroindole on the expression of *Th* and *Vmat2* at 24 h are concentration dependent. All data are the result of three or more independent measurements and are expressed as an average ± standard deviation (SD), with *p<0.05, **p<0.01, ***p<0.001, ****p<0.0001 by one-way ANOVA, in comparison with the response for vehicle treatment (DMSO).

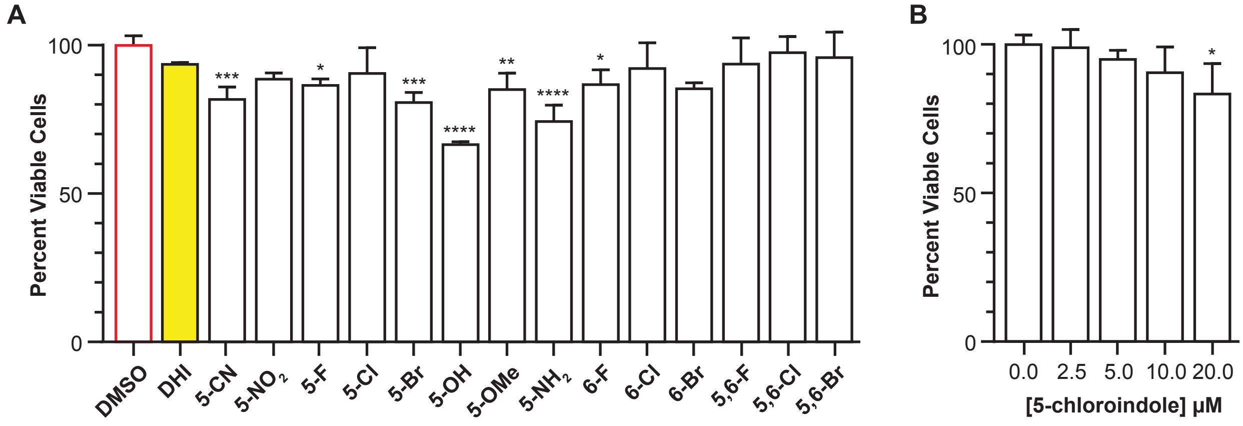


**Supplemental Figure 2C.** 5-chloroindole is not cytotoxic. **(A)** Approximately half of the indoles tested reduce the percentage of viable of MN9D cells following treatment with 10 μM compound for 24 h. **(B)** 5-chloroindole has no significant effect on cell viability at concentrations ≤ 10 μM following treatment for 24 h. Cell viability was measured using CytoTox-Glo Cytotoxicity Assay Kit (Promega) according to the manufacturer's instructions following treatment of cells (10,000 cells/well) with the indicated indole or DMSO. All experimental values are the result of three independent measurements ± SD, with *$p<0.05$, **$p<0.01$, ***$p<0.001$, ****$p<0.0001$ by one-way ANOVA, in comparison with the response for vehicle treatment (DMSO).

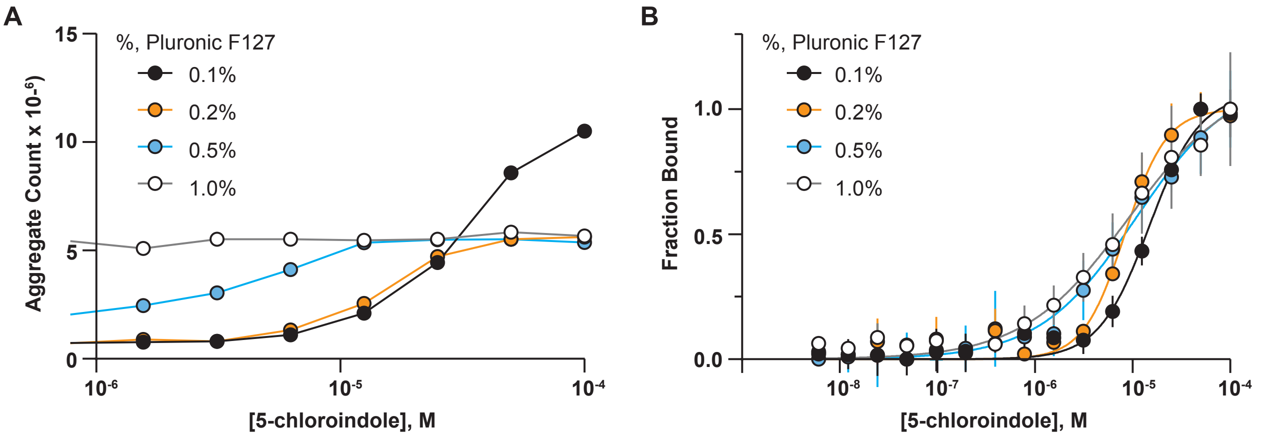


**Supplemental Figure 3A.** Increasing concentrations of surfactant decrease the formation of 5-chloroindole nanoaggregates and increase the affinity for the Nurr1 LBD by less than two-fold. **(A)** Aggregate count (DLS normalized intensity) for 5-chloroindole measured with increasing percentages of Pluronic F127. **(B)** Binding affinity of 5-chloroindole measured with increasing percentages of Pluronic F127; $K_D$ (0.1%) = 15.0 ± 1.2 μM, ($n_H$ = 2); $K_D$ (0.2%) = 8.3 ± 0.7 μM, ($n_H$ = 2); $K_D$ (0.5%) = 10.9 ± 0.3 μM, ($n_H$ = 1); $K_D$ (1.0%) = 9.1 ± 0.4 μM, ($n_H$ = 1). All experimental values are the result of three or more independent biological replicates ± standard deviation.

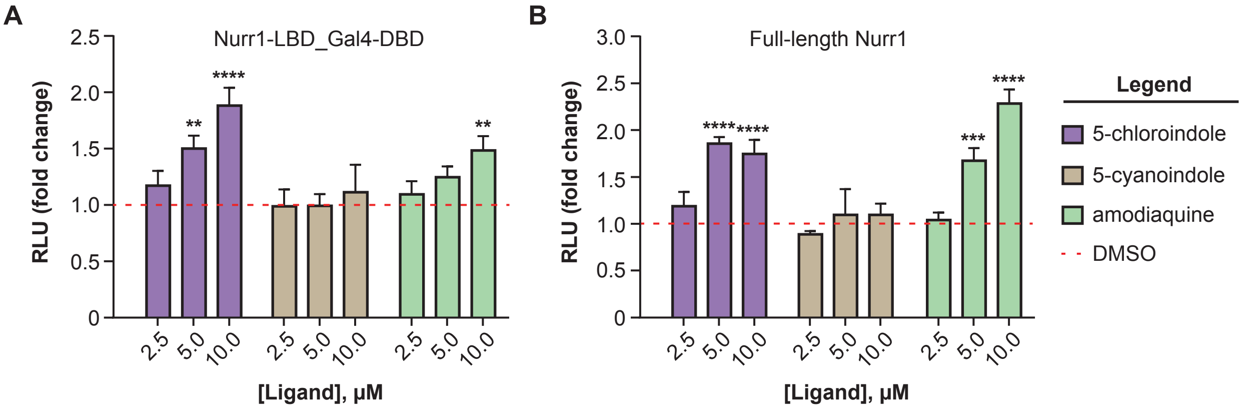


**Supplemental Figure 3B.** The DHI analog 5-chloroindole stimulates Nurr1 activity in two different luciferase reporter assays. In both the **(A)** Nurr1-LBD_Gal4-DBD luciferase reporter assay and the **(B)** full-length Nurr1 NBRE luciferase reporter assay, 5-chloroindole stimulates production of luciferase. Control compounds 5-cyanoindole (negative control) and amodiaquine (positive control) perform as expected. MN9D cells were individually treated with the indicated concentrations of ligands for 6 h prior to measuring luciferase signal (RLU, relative luminometer units; see Supporting Information for additional details). All experimental values are the result of three or more independent biological replicates and are expressed as the relative average response ± standard deviation, with *$p<0.05$, **$p<0.01$, ***$p<0.001$, ****$p<0.0001$ by one-way ANOVA, in comparison to the response with vehicle (DMSO) only.

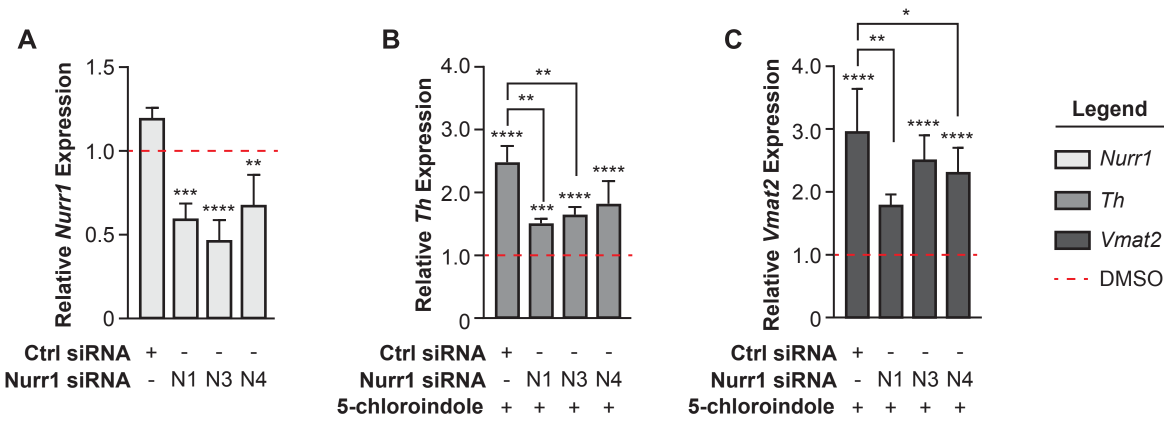


**Supplemental Figure 3C** The effect of 5-chloroindole on the expression of Nurr1 target genes depends on the expression of Nurr1. Expression of *Nurr1*, *Th* and *Vmat2* transcripts was determined in the presence of 5-chloroindole (10 μM, 24 h) with (Nurr1 siRNA) or without (Ctrl siRNA) knockdown of *Nurr1* levels. Gene expression levels in the presence of 5-chloroindole are relative to the same treatments with vehicle (DMSO) only. **(A)** The expression of Nurr1 is significantly reduced by Nurr1 siRNA, but not control siRNA. Knockdown of *Nurr1* in MN9D cells expressing endogenous Nurr1 with Nurr1 siRNA was carried out as described in Supporting Information. **(B, C)** The effect of 5-chloroindole on the expression of *Th* and *Vmat2* is significantly reduced in the presence of Nurr1 siRNA, but not control siRNA. All experimental values are the result of three or more independent biological replicates and are expressed as the relative average response ± standard deviation, with * $p<0.05$, ** $p<0.01$, *** $p<0.001$, **** $p<0.0001$ by one-way ANOVA, in comparison to the response with vehicle (DMSO) only.

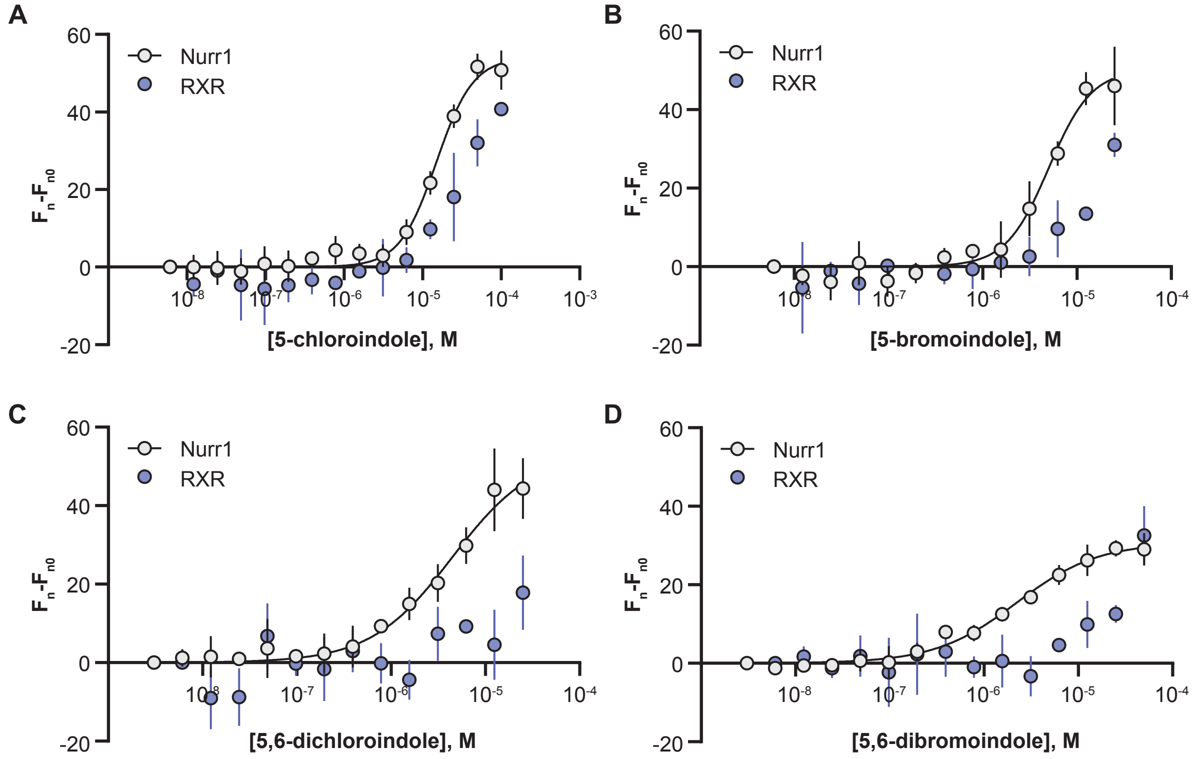


**Supplemental Figure 3D.** Halogenated indoles bind specifically to the Nurr1 LBD, but not the RXRα LBD. Comparison of the MST binding isotherms for **(A)** 5-bromoindole, **(B)** 5-chloroindole, **(C)** 5,6-dibromoindole, and **(D)** 5,6-dichloroindole reveal saturable binding to the Nurr1 LBD (grey circles), but not to the RXRα LBD (blue circles). Binding assays were carried out as described in Supporting information. All experimental values are the result of three or more independent biological replicates ± standard deviation.

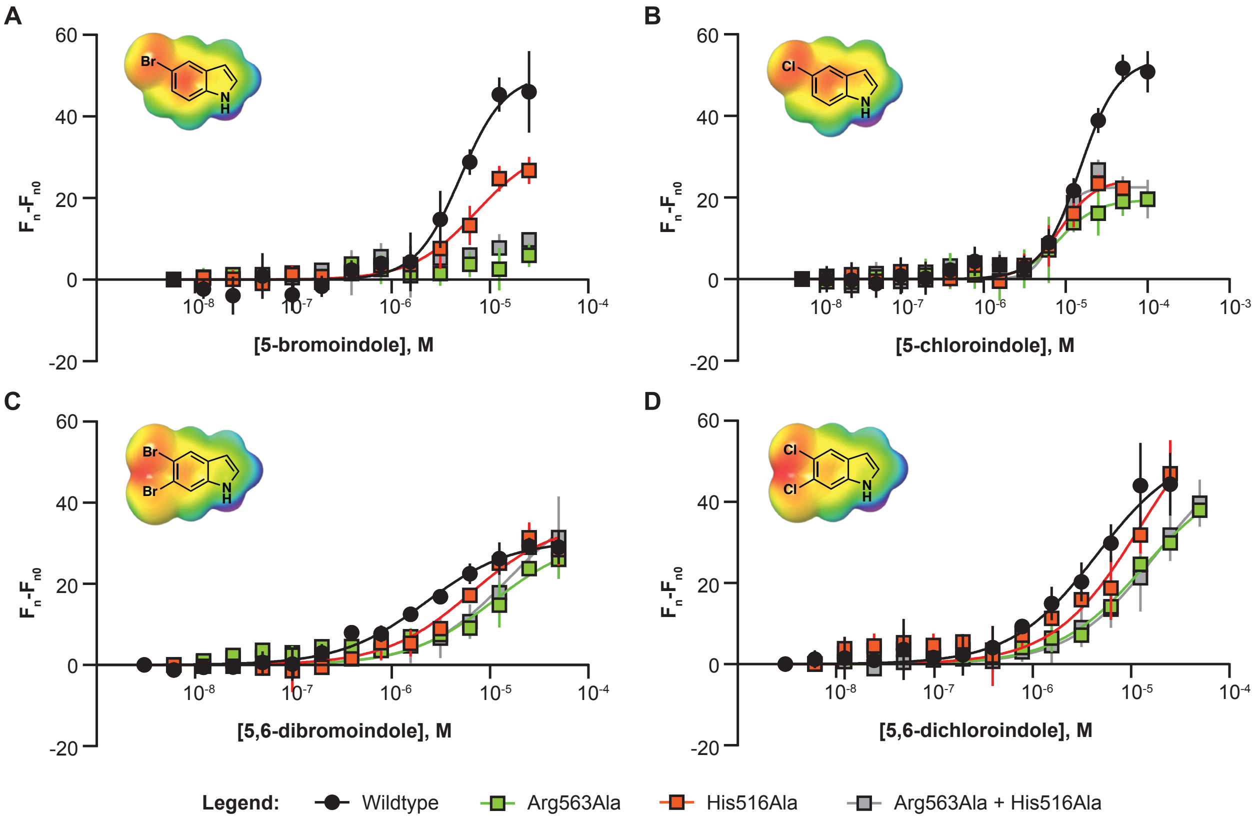


**E**

| Compound | Nurr1 Variant | Affinity ($K_D$, μM) | Response Amplitude |
|---|---|---|---|
| 5-bromoindole | Wildtype | 5.0 ± 0.7 | 50 ± 4 |
| | Arg563Ala | ND | ≤10 (****) |
| | His516Ala | 7.3 ± 2.1 | 33 ± 5 (*) |
| | Arg563Ala + His516Ala | ND | ≤10 (****) |
| 5-chloroindole | Wildtype | 15.0 ± 1.2 | 54 ± 2 |
| | Arg563Ala | 8.3 ± 1.7 | 19 ± 2 (****) |
| | His516Ala | 8.6 ± 1.2 | 25 ± 2 (****) |
| | Arg563Ala + His516Ala | 7.3 ± 0.8 | 23 ± 1 (****) |
| 5,6-dibromoindole | Wildtype | 2.2 ± 0.2 | 31 ± 1 |
| | Arg563Ala | 11.0 ± 2.8 | 32 ± 3 (ns) |
| | His516Ala | 6.4 ± 1.3 | 35 ± 2 (ns) |
| | Arg563Ala + His516Ala | 15.6 ± 4.3 | 43 ± 5 (ns) |
| 5,6-dichloroindole | Wildtype | 4.5 ± 0.9 | 54 ± 4 |
| | Arg563Ala | 13.1 ± 2.2 | 47 ± 3 (ns) |
| | His516Ala | 12.0 ± 3.6 | 67 ± 9 (ns) |
| | Arg563Ala + His516Ala | 17.9 ± 3.8 | 54 ± 5 (ns) |

**Supplemental Figure 4.** Mutation of residues (His516, Arg563) within the “566 site” significantly impacts the binding of monosubstituted indoles, but not the corresponding disubstituted indoles. **(A, B)** Single and double mutants *increase* the affinity of the 5-substituted indoles for the receptor and dramatically alter the thermophoresis response amplitude. **(C, D)** Single and double mutants *decrease* the affinity of disubstituted indoles for Nurr1, but have relatively small effects on the thermophoresis response amplitude. **(E)** Table summarizing the binding data values ($K_D$, MST amplitude) for the graphs shown in A-D. All experimental values are the result of three or more independent biological replicates ± standard deviation.

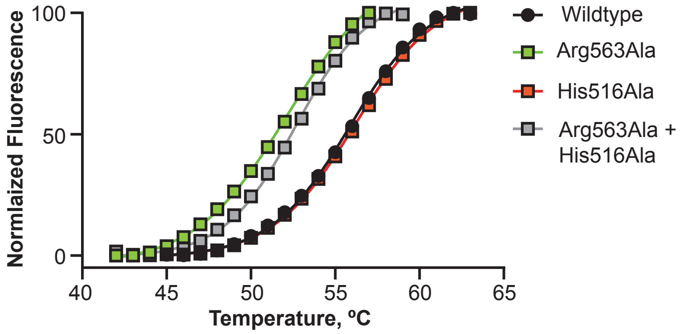


| Nurr1 Variant | $T_m$ ºC ± SEM |
|---|---|
| Wildtype | 55.80 ± 0.05 |
| Arg563Ala | 52.11 ± 0.07 |
| His516Ala | 56.16 ± 0.06 |
| Arg563Ala + His516Ala | 52.64 ± 0.05 |

**Supplemental Figure 5.** Mutation of Arg563 within the Nurr1 LBD reduces the thermal stability of the protein. Melting curves were acquired using differential scanning fluorimetry (DSF). The Nurr1 LBD (4 μM), dissolved in 25 mM HEPES buffer, pH 7.4, 150 mM NaCl, 1xSYPRO$^{TM}$ Orange dye. The fluorescence response was normalized to the largest fluorescent value, defined as 100%, within each data set. The reported Tm (the inflection point of the sigmoidal curve) was calculated using the Boltzmann sigmoid equation: Y = bottom + (top-bottom)/(1 +exp((Tm-x/slope)), where bottom and top are the values of the minimum and maximum intensities.  Each data point is the average of at least three independent measurements ± standard deviation; the curve for each variant is the result of the global fit to all replicates.

### EXPERIMENTAL PROTOCOLS

#### Chemicals and Reagents

The indoles used in the present study were purchased from Ambeed or Fisher Scientific. All other chemicals were purchased from Millipore, Sigma, or ThermoFisher Scientific, unless otherwise indicated. The pYFJ16-LplA(W37V) plasmid used to express the coumarin ligase (LplA) was purchased from Addgene. The MN9D Tet-ON cell line was graciously provided by Dr. Thomas Perlmann (Karolinska Institute). The reporter plasmid NBREx3-POMC-Luc was graciously provided by Dr. Jacques Drouin (Institut de Recherches Cliniques de Montréal, Canada).

#### Computational Methods

The molecular electrostatic potential surface for each indole was calculated using the 6-31G** basis sets and the B3LYP-D3 functional in water (PBS solvent model), and bromine atoms were treated with the LAV2P**. All calculations were performed using Jaguar (Schrodinger®) software.

Models for non-covalent binding of substituted-indoles to Nurr1 within the DHI-binding 566 site were prepared according to quantum mechanics-molecular mechanics (QM/MM) calculations using the dispersion-corrected (D3) Density Functional Theory (DFT) and the LACVP* basis sets for the QM region, and the MM OPLS2005 force field for the other residues. Qsite (Schrodinger®) was used for these calculations.

Single point interaction energies between the side chain of His516 and the C5-substituted indoles were calculated using Jaguar (Schrodinger®) with the LMP2/cc-pVDZ** level of theory. Energies were calculated for the gas phase, rather than with implicit solvation models, as nuclear receptor ligand binding pockets are traditionally hydrophobic cavities; this is certainly the case for the previously identified DHI-binding 566 site (PDB:1OVL). The coordinates of the complexes used for these calculations were taken from QM/MM optimized, non-covalently bound indole-Nurr1 structures at the DFT-D3/LACVP* level of theory. All energy values were calculated in kcal $mol^{-1}$.

The pKa predictions for His516 were made using by propKa 3.1 after QM/MM optimization of the non-covalent ligand-bound Nurr1 structures. In the starting structure (PDB ID: 6DDA), the predicted pKa for His516 is 6.5. Upon ligand binding and optimization, the pKa value of His516 is increased, especially for substituted-indoles containing hydrogen bond acceptors at C-5 position.

#### DNA Constructs

The plasmid used for the expression of LAP2-tagged Nurr1 LBDs (LAP2 Nurr1) was prepared by GenScript (Piscataway, NJ) as the product of gene synthesis and subcloning into pET-21a(+) Vector (GenScript) using the NdeI and XhoI sites within the MCS. The protein sequence of the resulting protein is shown in **Supplementary Table 1**. Mutants of LAP2 Nurr1 were prepared by GenScript starting from LAP2 Nurr1 vector.

**Supplementary Table 1**

| Construct Name | Sequence |
|---|---|
| **LAP2 Nurr1** | MKKGHHHHHHGFEIDKVWYDLDAGAISLISALVRAHVDSNPAMTSLDY SRFQANPDYQMSGDDTQHIQQFYDLLTGSMEIIRGWAEKIPGFADLPK ADQDLLFESAFLELFVLRLAYRSNPVEGKLIFCNGVVLHRLQCVRGFG EWIDSIVEFSSNLQNMNIDISAFSCIAALAMVTERHGLKEPKRVEELQN KIVNCLKDHVTFNNGGLNRPNYLSKLLGKLPELRTLCTQGLQRIFYLKL EDLVPPPAIIDKLFLDTLPF |
| **LAP2 RXRα** | MKKGHHHHHHGSGSENLYFQSGSGSGSGFEIDKVWYDLDAGSGSDMP VERILEAELAVEPKTETYVEANMGLNPSSPNDPVTNICQAADKQLFTLV EWAKRIPHFSELPLDDQVILLRAGWNELLIASFSHRSIAVKDGILLATGL HVHRNSAHSAGVGAIFDRVLTELVSKMRDMQMDKTELGCLRAIVLFNP DSKGLSNPAEVEALREKVYASLEAYCKHKYPEQPGRFAKLLLRLPALR SIGLKCLEHLFFFKLIGDTPIDTFLMEMLEAPHQMT |

**Synthesis and Purification of the Azide-reactive Fluorescein Probe for MST**

The dibenzocyclooctyne (DBCO)-5/6-carboxyfluorescein probe was synthesized according to previously reported procedures (Patent US20150125904A1). Briefly, dibenzocyclooctyne-amine (3.2 mg, 11.5 μmol) in 540 μL anhydrous DMF was added to 5/6-carboxyfluorescein *N*-succinimidyl ester (6 mg, 12.7 μmol) and triethylamine (4.9 μL, 35.7 μmol). After stirring overnight at ambient temperature, the solvent was removed by lyophilization and resulting oil was resuspended in EtOAc and extracted against 1 M HCl, followed by saturated NaCl. The organic layer was dried over anhydrous $MgSO_4$ and then concentrated to dryness via rotary evaporation to give the crude alkyne. The desired product was purified to homogeneity using preparative thin layer chromatography (EtOAc), eluted from the silica gel (EtOAc:MeOH, 95:5), and then concentrated to dryness to give the final product, dibenzocyclooctyne (DBCO)-5/6-carboxyfluorescein, in an overall yield of 60%. ESI-MS characterization $[M+H]^+$ gave 635.7 observed; 635.2 calculated.

**Protein Expression and Purification**

The Nurr1 LBD protein, containing the N-terminal "LAP2" sequence that is recognized by a "coumarin ligase", was expressed and purified using metal affinity and size-exclusion chromatography according to the previously reported protocol, except that the TEV cleavage step and reverse metal affinity chromatography were omitted.[1] The RXRα LBD protein, containing the N-terminal "LAP2" sequence, was expressed and purified identically to the Nurr1 LBD protein with minor modifications. Specifically, the elution of protein from Talon resin was performed by a step gradient of 10 CV each of 50 mM, 100 mM, 200 mM and 300 mM imidazole in 50 mM Tris-HCl buffer containing 300 mM NaCl at pH 7.8. Purity of protein in each fraction was analyzed by SDS PAGE and fractions eluted with 100-300 mM imidazole were pooled and concentrated. Resulting protein solution was then applied to a S75 10/300 SEC column (GE Healthcare Life Sciences) using a running buffer composed of 50 mM Tris-HCl, 100 mM KCl, 1 mM DTT, 10 % glycerol at pH 8.0. The sequence of the construct (prepared by GenScript) is shown above in Supplementary Table 1.

**Labeling Protein with Fluorescein for MST Assays**

The fluorescein probe was ligated to the N-terminal LAP2 tag within Nurr1 LBD and RXR LBD using a re-engineered version of the enzyme lipoic acid ligase (LplA) from Escherichia coli as previously reported.[2, 3] Briefly, the plasmid harboring the gene coding for “coumarin ligase” [pYFJ16-LplA(W37V); Addgene] was transformed into BL21(DE3) cells (New England BioLabs) and a single colony was subsequently used to inoculate LB media supplemented with 100 µg/mL ampicillin, and the culture was grown at 37 ºC until reaching an $OD_{600}$ of 0.9, at which point protein expression was induced by adding IPTG (100 µM final concentration) and incubation continued for 16 hours at 25 ºC. Next, the cells were harvested by centrifugation (3,500 g, 20 minutes, 4 ºC) and the pellet was resuspended in lysis buffer (50 mM Tris base, 300 mM NaCl, pH 7.8) containing cOmplete mini EDTA-free protease inhibitor cocktail (Roche). Cells were lysed by continuous passage at 15,000 psi using C3 Emulsiflex (Avestin). The extract was cleared by centrifugation (21,000 g, 45 minutes, 4 ºC) and the $His_6$-tagged enzyme was purified using Ni-NTA agarose (Qiagen). Fractions were analyzed by 12% SDS-PAGE followed by Coomassie staining. Fractions containing LplA were pooled and dialyzed against 20 mM HEPES, 150 mM NaCl, 1 mM DTT, 10 % glycerol, pH 8.0. The protein concentration was determined by measuring the A280 and using the calculated extinction coefficient 41,940 $M^{-1}$ $cm^{-1}$.

Sequence specific (LAP2-tag) incorporation of the fluorescein probe was achieved according to the previously reported protocol.[4] A typical reaction contained LAP2-tagged protein (20 µM), buffer (25 mM sodium phosphate, pH 7.0, 2 mM magnesium acetate, 1 mM ATP),10-azadecanoic acid (100 µM), and the enzyme W37V LplA (1 µM). After incubating at 30°C for 1 h, the reaction was supplemented with 200 µM DBCO-linked 5(6)-carboxyfluorescein probe and allowed to incubate at ambient temperature for 30 min before being buffer exchanged into 25 mM HEPES, pH 7.4, 150 mM NaCl. The concentration of fluorescent label was determined by UV-vis spectroscopy using the extinction coefficient $\varepsilon_{493}$ = 70,000 $M^{-1}cm^{-1}$ for fluorescein.

**Microscale Thermophoresis Assay**

Concentration-dependent association of the indoles with the Nurr1 LBD was carried out using microscale thermophoresis. Stock solutions (10 mM in DMSO) of each indole were serially diluted (200 µM indole, 0.5x dilutions down to 0.0061 µM) in MST buffer (25 mM HEPES, pH 7.4, 150 mM NaCl, and 0.1% Pluronic F127) containing 2% DMSO. The dilutions were carried out with 4% DMSO in the MST buffer for a total of 16 concentrations. Equivalent volumes and concentrations of the fluorescently labeled Nurr1 LBD in MST buffer were added to each ligand dilution in the series to reach a final concentration of 75 nM labeled protein. After incubating for 20 minutes, the samples were loaded into Monolith NT.115 Capillaries (Nanotemper).

Data were collected using the Monolith NT.115 System (Nanotemper), with settings for all samples at 40% excitation power and 40% MST power. The initial fluorescence was recorded for 3 sec and the thermophoresis fluorescence response was recorded for 20 sec. The data was inspected with Palmist software[5], and data points affected by initial fluorescence quenching or photobleaching were eliminated. The fluorescent response for each sample was normalized to the initial fluorescence using Palmist software to provide the values of thermophoresis ($F_n$). The resulting data was used to generate a plot of the change in thermophoresis ($F_n$-$F_{n0}$, where $F_n$ = thermophoresis and $F_{n0}$ = thermophoresis response in unbound range) versus concentration of the ligand. GraphPad Prism v. 8.3.0 software was used to fit the resulting data to a mass action equation for a specific binding with Hill slope, solving for $K_D$: $F_n$-$F_{n0}$ = $F_{max}\cdot[L]^n/(K_D^n+[L]^n)$, where $F_{max}$ = maximum amplitude of thermophoresis, [L] = concentration of the ligand at a specific point, $K_D$ = dissociation constant, $n_H$ = Hill Coefficient.

MST binding assays for the 5-chloroindole, 5-bromoindole, 5,6-dichloroindole, and 5,6-dibromoindole were also run using an unrelated protein, the RXR$\alpha$ LBD, and demonstrate that the signal changes observed for binding to the Nurr1 LBD are not dominated by non-specific binding artifacts (**Supplemental Figure 3B**).To investigate the potential impact of compound nanoaggregation on binding affinity, we used dynamic light scattering to inspect the aggregation of 5-chlorindole in solution and repeated the MST binding experiments in the presence of increasing concentrations of the surfactant Pluoronic F127 (**Supplemental Figure 3D**). The nanoaggregation properties of 5-chloroindole were found to be minimal, ruling out the possibility that the observed concentration-dependent changes in the MST signal are dominated by compound aggregation. As well, increasing concentrations of the surfactant Pluronic F127 had small effects (<2-fold) on the affinity of 5-chloroindole for the Nurr1 LBD (**Supplemental Figure 3D**). Furthermore, UV/VIS spectroscopy ($A_{280}$) of both 5-chloro- and 5-bromoindole, under conditions equivalent to those used in the MST binding assays (0.1% Pluronic F127), reveals that the absorbance of both compounds remains linear over all concentrations tested, indicating that these compounds do not precipitate under the assay conditions (**data not shown**).

**Differential Scanning Fluorimetry Assay**

The Nurr1 LBD protein was buffer exchanged into 25 mM HEPES, 150 mM NaCl, pH 7.4 using a Zeba Spin Desalting Column (ThermoFisher). The DSF assays were carried out in a final volume of 30 µL, comprised of 4 mM protein, 1x SYPRO Orange (ThermoFisher/Life Technologies, from 5000x stock), and buffer comprised of 25 mM HEPES, pH 7.4, 150 mM NaCl. Samples were allowed to incubate in the dark for 30 min at 25 °C, prior to exposure to thermal gradient. Fluorescence was monitored using the CFX Connect Real-Time PCR Detection System (BioRad) in a 96-well plate (BioRad). The thermal gradient was executed from 25 °C to 95 °C at a rate of 0.05 °C/s. The fluorescence response was normalized so that 0% and 100% are defined as the smallest and the largest mean in each dataset, correspondingly. Melting temperatures, $T_m$ (the inflection point of the sigmoidal curve), was calculated using the Boltzmann sigmoid equation: Y = bottom + (top-bottom)/(1 +exp($T_m$-x/slope)), where bottom and top are the values of the minimum and maximum intensities.

**Dynamic Light Scattering Assay**

5-Chloroindole was serially diluted from 10 mM DMSO stock into MST buffer supplemented with various amounts of Pluronic F127 (0.1 %, 0.2 %, 0.5 %, 1.0 %) at room temperature for a final concentration of 0.2% DMSO. Measurements were made using a DynaPro MS/X (Wyatt Technology) with a 55 mW laser at 826.6 nm, using a detector angle of 90°. The laser power was 100%, and the acquisition time was 2 s. Histograms represent the average of three independent data sets, each with at least 10 measurements.

**Cellular Assays**

*Target Gene Transcription Assays*. These assays were carried out using standard protocols. MN9D TET-ON frozen cell stocks (P5) were thawed and grown for 60-72 h on poly-D-lysine pre-treated culture dishes (100 mm) in Dulbecco's Modified Eagle Medium/Nutrient Mixture F-12 (DMEM/F-12; Gibco) supplemented with 5% Tet System Approved FBS (Takara

Bio USA) at 37°C, 5% $CO_2$ to ~80 % confluency. The resulting cells were then trypsinized with 0.25% Trypsin-EDTA (Gibko) and diluted to $2 \cdot 10^5$ cells/ml with fresh medium. The resulting cell suspension (0.8 mL) was added to 2x concentrated compound in the same medium containing 0.2 % DMSO (0.8 mL) in an Eppendorf tube. The cell suspension with compound or vehicle (DMSO) was then seeded onto a 24-well plate pre-treated with Poly-D-Lysine at 0.8 mL per well. Assays were performed under conditions of basal Nurr1 expression, *without* induction of additional Nurr1 expression using doxycycline.

After 24 h, total RNA was extracted the cells in each well using the Quick-RNA MiniPrep Plus Kit (Zymo Research), according to the manufacturer's instructions. The cDNA was then synthesized from 1000 ng of purified RNA using High-Capacity cDNA Reverse Transcription Kit (Applied Biosystems) and used as template. The qPCR as performed using iTaq Universal SYBR Green Supermix (BioRad) and CFX96 Real-Time Detection System machine (BioRad). Briefly, qPCR was performed in hard-shell 96-well PCR plates (BioRad) using cDNA corresponding to 8.75 ng of starting total RNA in a volume of 15 μl, containing 7.5 μl of SYBR Green Supermix, and 1 μL of 10 μM forward and reverse primers. Cycling parameters for qPCR included an initial denaturation at 95°C for 3 min, followed by 40 cycles of 95°C for 5 s and annealing at 56°C for 30 s. The forward and reverse primers (**Supplementary Table 2**) were ordered from IDT. Gene expression was quantified by the comparative2-ΔΔCt method, with the mouse housekeeping gene hypoxanthine-guanine phosphoribosyltransferase (*Hprt*) used as internal reference to determine the relative mRNA expression. Transcript levels for target genes were normalized to the housekeeping gene *Hprt* and fold change was compared to gene expression levels from vehicle (DMSO) only treated cells. GraphPad Prism 8 software was used for statistical analysis. Two-way ANOVA was applied for DMSO fold change vs compound. Results are from three independent experiments. Relative average expression ± SD; *$p < 0.05$, **$p < 0.01$, ***$p < 0.001$, ****$p < 0.0001$ by ANOVA in comparison expression with 0 μM compound (DMSO only).

**Supplementary Table 2.** Sequences of primers used in RT-qPCR

| Target Gene | Primer | Sequence (5′ to 3′) | Reference |
|---|---|---|---|
| ***Hprt*** | FW | TGGGAGGCCATCACATTGT | Volpicelli, Floriana, et al. "Direct regulation of Pitx3 expression by Nurr1 in culture and in developing mouse midbrain." *PloS one* 7.2 (2012): e30661. |
| | REV | AATCCAGCAGGTCAGCAAAGA | |
| ***Nurr1*** | FW | CAACTACAGCACAGGCTA | |
| | REV | GCATCTGAATGTCTTCTACCTTAATG | |
| ***Th*** | FW | TCCAACCTTTCCTGGCCCAG | Hwang, Dong-Youn, et al. "Vesicular monoamine transporter 2 and dopamine transporter are molecular targets of Pitx3 in the ventral midbrain dopamine neurons." *Journal of neurochemistry* 111.5 (2009): 1202-1212. |
| | REV | GCATGAAGGGCAGGAGGAAT | |
| ***Vmat2*** | FW | GAAGTCCACCTGCTAAGGAAGAA | Designed in this work. |
| | REV | TCACTGGAGACACATGTACACAG | |

Nurr1 knockdown (siRNA) assays were completed according to standard protocols. Briefly, MN9D cells were resuspended in DMEM/F-12 with 5% FBS at $1 \cdot 10^6$ cells/mL, and reverse-transfected with either control or Nurr1 siRNAs (40 nM final concentration) by combining 4 mL of cell suspension with 1 mL of Opti-MEM containing 200 nM siRNA and 10 µL Lipofectamine 2000 (Invitrogen, cat#:11668019). The resulting cell suspension was plated on poly-D-lysine treated six-well plates at 2.5 mL/well, and allowed to incubate for 24 h at 37 °C, in 5% $CO_2$ incubator. After 24 hours, the cells were trypsinized, resuspended in DMEM/F-12 with 5% FBS at $2 \cdot 10^5$ cells/mL, mixed with equal volume of 20 µM 5-chloroindole in the same media containing 0.2% DMSO (prepared by diluting DMSO stock of 5-chloroindole (10 mM) in warm DMEM/F-12 with 5% FBS; for the DMSO control an equivalent volume of DMSO was used instead of the compound stock), and immediately re-plated on poly-D-lysine pre-treated 24-well plates at $1 \cdot 10^5$ cells/well and incubated as above. After 24 hours, total RNA was isolated and the target gene transcription assay (qPCR) was performed as described above. Nurr1 siRNA N1 were purchased from Sigma (Cat No. SASI_Mm02_00322368), and the sequences were as follows: 5'- GAA UCA GCU UUC UUA GAA U[dT][dT] -3' (sense) and 5'-AUU CUA AGA AAG CUG AUU C[dT][dT] -3' (antisense). Nurr1 siRNA N3 and N4 were ordered from IDT, and the sequences were as follows: 5'-GCAUCGCAGUUGCUUGACATT ( N3 sense) and 5'-UGUCAAGCAACUGCGAUGCGT (N3 antisense); 5'-CUAGGUUGAAGAUGUUAUAGGCACT (N4 sense) and 5' AGUGCCUAUAACAUCUUCAACCUAGAA (N4 antisense).[6]
GFP negative control DsiRNA were purchased from IDT (Cat No. 51-01-05-06).

Cytotoxicity assays were carried out using the CytoTox-Glo Cytotoxicity Assay Kit (Promega) according to the manufacturer's instructions. Briefly, MN9D cells were resuspended in DMEM/F-12 with 5% FBS at $2 \cdot 10^5$ cells/ml and added to an equal volume of two-fold concentrated compound in the same media containing 0.2% DMSO (prepared by diluting DMSO stock of the compound in warm DMEM/F-12 with 5% FBS; for DMSO control equivalent volume of DMSO was used instead of the compound stock), and immediately re-plated onto poly-D-lysine pre-treated 96-well white-walled flat clear bottom plate (Corning cat#3903) at a density of $1 \cdot 10^4$ cells/well (100 µL/well), 3 replicas per compound. As a background (BG) control, for each of the compound (or DMSO) equivalent volume of compound mixed with DMEM/F-12 + 5% FBS was plated. After 24 h treatment at 37 °C, in 5% $CO_2$ incubator, 50 µL of CytoTox-Glo™ Cytotoxicity Assay Reagent was added to each well, mixed briefly by orbital shaking and incubated for 15 min at ambient temperature. Luminescence signal corresponding to dead cells was measured using Biotek Synergy H4 hybdrid microplate reader. After the measurement, 50 µL of Lysis Reagent was added to each well, mixed, incubated for 15 min at ambient temperature, and total luminescence was measured. The percentage of viable cells was then calculated as follows: Percent Viable Cells (%) = 100%·(Total cell luminescence (test compound) – Dead cell luminescence (test compound))/(Total cell luminescence (DMSO) – Dead cell luminescence (DMSO)).

*Luciferase Reporter Assays*. These assays were executed using standard protocols. MN9D TET-ON cells (P5) were grown for 60-72 h (see above) before being seeded at $1 \cdot 10^5$ cell/well in 24-well plate in DMEM/F12 + 5% TET-ON approved FBS 3 h before transfection. Cells were transfected using Lipofectamine 2000 (Invitrogen) and plasmid DNAs according to manufacturer's instructions. Lipofectamine/DNA complexes were prepared in Opti-Mem medium (Gibco) and incubated with cells overnight. The Nurr1-LBD_Gal4-DBD expressing plasmid was prepared by subcloning the Nurr1 LBD fragment into pM plasmid (Clontech) containing GAL4 DBD. The reporter plasmid pGL4.35 (luc2P/9XGAL4 UAS/Hygro) Vector (Promega) contains nine repeats of GAL4 UAS (Upstream Activator Sequence) and drives transcription of the luciferase reporter gene luc2P in response to binding of Nurr-LBD_Gal4-DBD chimeric protein. The pRL-null (Promega) plasmid expressing renilla luciferase was used as an internal control. The amounts of pM-Nurr1-LBD_Gal4-DBD, pGL4.35, pRL-null and Lipofectamine 2000 used for transfections were 100 ng, 100 ng, 200 ng and 1 μL per well respectively. Alternatively, cells were co-transfected with the reporter plasmid NBREx3-POMC-Luc containing three copies of the NBRE sequence (5′-GATCCTCGTGCGA**AAAGGTCA**AGCGCTA-3′) subcloned into the pXP1-luc plasmid containing the minimal (positions -34 to +63) POMC promoter as described previously[7], and the pRL-null plasmid. The amounts of NBREx3-POMC-Luc, pRL-null and Lipofectamine 2000 used for transfecting cells were 100 ng, 200 ng and 1 μL per well respectively.

Transfected cells were treated with increasing concentrations of 5-chloroindole or vehicle only for 6 h, after which time the media was aspirated and luciferase activity was measured. Cells from each well were incubated for 15 min with 220 μL/well of Dual-Glo® Luciferase Reagent (Promega) at room temperature upon rotation on the orbital shaker. Resulting lysates were cleared from the cell debris by centrifugation for 2 min at 16,000 rcf. Resulting solutions were transferred to a white opaque 96-well plate (65 μL/well; 3 wells/sample) and firefly luciferase activity was measured using Veritas Microplate Luminometer (Turner BioSystems, Sunnyvale, CA). An equal volume of Dual-Glo® Stop & Glo® Reagent (Promega) was added to each well. Renilla luciferase activity was measured after 20 min of incubation of the plate inside the luminometer. Experimental values are expressed as the average of firefly/renilla luciferase activity ± standard deviation (for three independent biological replicates). One-way Analysis of Variance (ANOVA) was used to determine statistical significance using GraphPad Prism 8.3.0 software, with *$p < 0.05$, **$p < 0.01$, ***$p < 0.001$, ****$p < 0.0001$ in comparison with 0 μM compound (DMSO only).
